# Symptom Dimension-Specific Neurotransmitter Correlates of Psychopathology and Cognition in Early Psychosis

**DOI:** 10.64898/2026.06.03.729892

**Authors:** Haley R. Wang, Charles H. Schleifer, Zhen-Qi Liu, Rachel A. McKinney, Rune Boen, Carolyn M. Amir, Hoki Fung, Bratislav Misic, Lucina Q. Uddin, Carrie E. Bearden, Katherine H. Karlsgodt

## Abstract

Extended duration of under-treated psychosis (DUP) is among the strongest predictors of poor outcome, yet diagnostic heterogeneity impedes treatment matching, with approximately 50% of patients failing to respond to first-line antipsychotics. Negative symptoms and cognitive impairment are particularly refractory, lacking effective pharmacological treatments. Identifying neurotransmitter systems associated with specific symptom dimensions could accelerate targeted therapeutic development and reduce DUP. We applied Partial Least Squares correlation (PLSc) to derive whole-brain resting-state functional connectivity (RSFC) and anatomical (cortical thickness and subcortical volume) signatures associated with five psychopathology dimensions (positive symptoms, negative symptoms, general psychopathology, mania, and cognition) in a transdiagnostic sample from the Human Connectome Project-Early Psychosis (HCP-EP; n=124). We tested associations with potential confounds including antipsychotic medication dosage and substance use. Signatures were spatially correlated with 21 Positron Emission Tomography (PET)-derived receptor and transporter maps across 9 neurotransmitter systems using the *neuromaps* toolbox. Significant RSFC signatures emerged for positive symptoms, negative symptoms, general psychopathology, and cognition, but not mania. The negative symptom RSFC signature correlated with norepinephrine transporter (NET; ρ=.40, *q*=.030) and vesicular acetylcholine transporter (VAChT; ρ=.38, *q*=.048) distributions. The cognition signature similarly correlated with VAChT (ρ=.48, *q*=.025). Anatomical signatures were associated with positive symptoms, general psychopathology, and cognition, but were more susceptible to confounding by medication and substance use. No significant receptor associations were detected for anatomical signatures. These findings implicate cholinergic and noradrenergic systems as molecular targets for negative symptoms and cognitive impairment, supporting prioritization of these systems in pharmacotherapy development in early psychosis.

## INTRODUCTION

Extended duration of under-treated psychosis (DUP) is among the strongest predictors of poor long-term outcome in psychosis spectrum disorders [1]. Reducing DUP, including by expediting identification of effective medications, remains a critical clinical priority [2]. One challenge in determining effective treatment is that diagnostic categories represent heterogeneous syndromes rather than biologically coherent entities [3, 4].

Consistent with this, medications in clinical practice are frequently prescribed to address specific symptoms that cut across diagnoses. However, regulatory guidance for development of psychopharmacological treatments for psychotic disorders mandates that pivotal trials enroll patients based on classification criteria for specific disorders rather than symptom profiles [5, 6], a framework that has been criticized for impeding translational research. This mismatch between diagnosis-based drug development and symptom-based clinical practice contributes to low treatment response rates, with approximately 50% of patients failing to respond to their first antipsychotic medication [7]. Trial-and-error prescribing further delays effective treatment. Certain symptom domains remain particularly refractory: negative symptoms show limited response to current antipsychotics [8, 9], and cognitive impairment, a core feature of psychosis strongly linked to functional outcome, lacks any FDA-approved pharmacological treatment [10–12]. Identifying precise molecular targets for different symptom domains could accelerate treatment matching and reduce DUP.

Decades of neurobiological research have implicated multiple neurotransmitter systems in psychosis, providing critical mechanistic insights, though investigations in humans have largely relied on diagnosis-based designs testing individual systems [13]. Dopamine dysregulation, particularly elevated presynaptic dopamine synthesis capacity in the striatum, is the most replicated finding and is a central mechanism of the majority of approved antipsychotics [14, 15]. In addition to dopamine, glutamatergic NMDA receptor hypofunction, serotonergic abnormalities, and GABAergic interneuron deficits have also been implicated in schizophrenia [14, 16]. The cholinergic system has received renewed attention given its putative role in cognitive deficits, culminating in the recent FDA approval of a muscarinic agonist for schizophrenia [17]. In addition, noradrenergic dysfunction has been linked to arousal, attention, and negative symptoms [18]. Despite this progress, most human studies have assessed one candidate neurotransmitter system at a time, typically comparing patients meeting diagnostic criteria for schizophrenia to unaffected controls [19]. This approach limits understanding of shared transdiagnostic biological underpinnings and precludes systematic comparison across neurotransmitter systems simultaneously.

To address these gaps, new methods are needed that can derive stable whole-brain neural signatures associated with specific symptom dimensions in a transdiagnostic framework and systematically characterize the molecular architecture of those brain patterns. Established multivariate methods make such an approach feasible. Partial Least Squares correlation (PLSc) is a multivariate technique that can be used to identify whole-brain patterns maximally covarying with clinical phenotypes, effectively translating clinical presentations into comprehensive neural signatures [20]. Prior work has demonstrated the utility of this approach in transdiagnostic early psychosis samples, revealing replicable brain signatures associated with symptom dimensions [21, 22]. Separately, the *neuromaps* toolbox aggregates PET-derived neurotransmitter receptor and transporter density maps from large healthy samples into a standardized atlas [23, 24]. Spatial correlation between brain signatures and receptor maps has been applied to characterize the molecular underpinnings of pharmacologically induced brain states and psychiatric conditions [25–28], enabling simultaneous examination of associations across all available receptor and transporter systems. This two-step approach, first translating symptoms into brain patterns and then annotating those patterns molecularly, could reveal candidate neurotransmitter systems for further mechanistic investigation and targeted therapeutic development.

In this study, we applied PLSc to derive whole-brain RSFC and anatomical signatures (patterns of functional connectivity and of regional cortical thickness and subcortical volume, respectively) associated with five dimensions of psychopathology (positive symptoms, negative symptoms, general psychopathology, mania, and cognition) in a transdiagnostic early psychosis (EP) cohort. As the first application of this approach in EP, we examined both neuroimaging modalities to determine which better captures dimension-specific patterns and receptor associations. We then correlated these signatures with 21 receptor and transporter maps spanning 9 neurotransmitter systems to identify dimension-specific brain signatures, characterize their receptor correlates, and evaluate convergence across modalities. We hypothesized that distinct symptom dimensions would be linked with distinct molecular associations that vary across neuroimaging modalities.

## METHODS

### Participants

The Human Connectome Project Early Psychosis Release 1.1 (HCP-EP) study cohort includes patients aged 16-35 years who were diagnosed with schizophrenia spectrum disorders, schizoaffective disorder, or psychotic mood disorders within three years of illness onset. The primary symptom analysis included 121 patients (mean age 22.7±3.7 years; 61.2% male) with usable resting-state fMRI and structural MRI data. The cognition analysis included a subset of 111 patients with available NIH Toolbox Cognition Battery data. Complete demographic and clinical characteristics are presented in **Table 1**.

**Table 1.** Demographic, clinical, and cognitive characteristics of the HCP-EP study participants.

| Characteristics | Symptom Analysis<br>(n=121) | Cognition Analysis<br>(n=110) |
| --- | --- | --- |
| Age, mean (SD), y | 22.7 (3.7) | 22.7 (3.6) |
| Male, No. (%) | 74 (61.2) | 68 (61.3) |
| Race, No. (%) |  |  |
| Asian | 8 (6.6) | 7 (6.3) |
| African American | 47 (38.8) | 45 (40.5) |
| American Indian/Alaska Native | 1 (0.8) | 1 (0.9) |
| White | 61 (50.4) | 54 (48.6) |
| Unknown or more than one race | 4 (3.3) | 4 (3.6) |
| Hispanic ethnicity, No. (%) | 8 (6.6) | 7 (6.3) |
| Parental education, mean (SD), y | 11.4 (2.5) | 11.4 (2.5) |
| WASI Full Scale IQ, mean (SD) | 101.6 (16.9) | 101.4 (16.4) |
| Duration of illness, mean (SD), y | 2.2 (1.4) | 2.2 (1.4) |
| SCID-5 diagnosis, No. (%) |  |  |
| Schizophrenia | 79 (65.3) | 76 (68.5) |
| Schizoaffective Disorder | 14 (11.6) | 10 (9.0) |
| Bipolar I or MDD with psychotic features | 28 (23.1) | 25 (22.5) |
| CPZ equivalent, mean (SD), mg | 157.4 (226.0) | 171.6 (230.8) |
| Cannabis use in past 30 days, No. (%) | 15 (12.4) | 15 (13.5) |
| PANSS scores, mean (SD) |  |  |
| Positive | 11.2 (3.9) | 11.0 (3.8) |
| Negative | 14.1 (5.4) | 14.2 (5.6) |
| General | 24.6 (5.3) | 24.7 (5.2) |
| YMRS score, mean (SD) | 4.8 (5.3) | 4.5 (5.2) |
| NIH Toolbox Cognition Composite Scores, mean (SD) |  |  |
| Total Composite | — | 100.7 (13.0) |
| Fluid Cognition Composite | — | 99.4 (14.3) |
| Crystallized Cognition Composite | — | 102.9 (10.2) |

**Note:** The cognition analysis sample (n=111) is a subset of the symptom analysis sample (n=121) comprising participants with available NIH Toolbox Cognition Battery data and usable neuroimaging data. The two samples did not differ significantly on any demographic or clinical variable (all $p>.05$ ). Abbreviations: WASI = Wechsler Abbreviated Scale of Intelligence; PANSS = Positive and Negative Syndrome Scale; YMRS = Young Mania Rating Scale; CPZ = chlorpromazine; SCID-5 = Structured Clinical Interview for DSM-5; MDD = Major Depressive Disorder. Schizophrenia includes schizophrenia, schizophreniform, psychosis NOS, delusional disorder, and brief psychotic disorder. Bipolar I or MDD with
psychotic features includes bipolar I disorder, bipolar II disorder, and major depressive disorder with psychotic features.

**Table 2.**
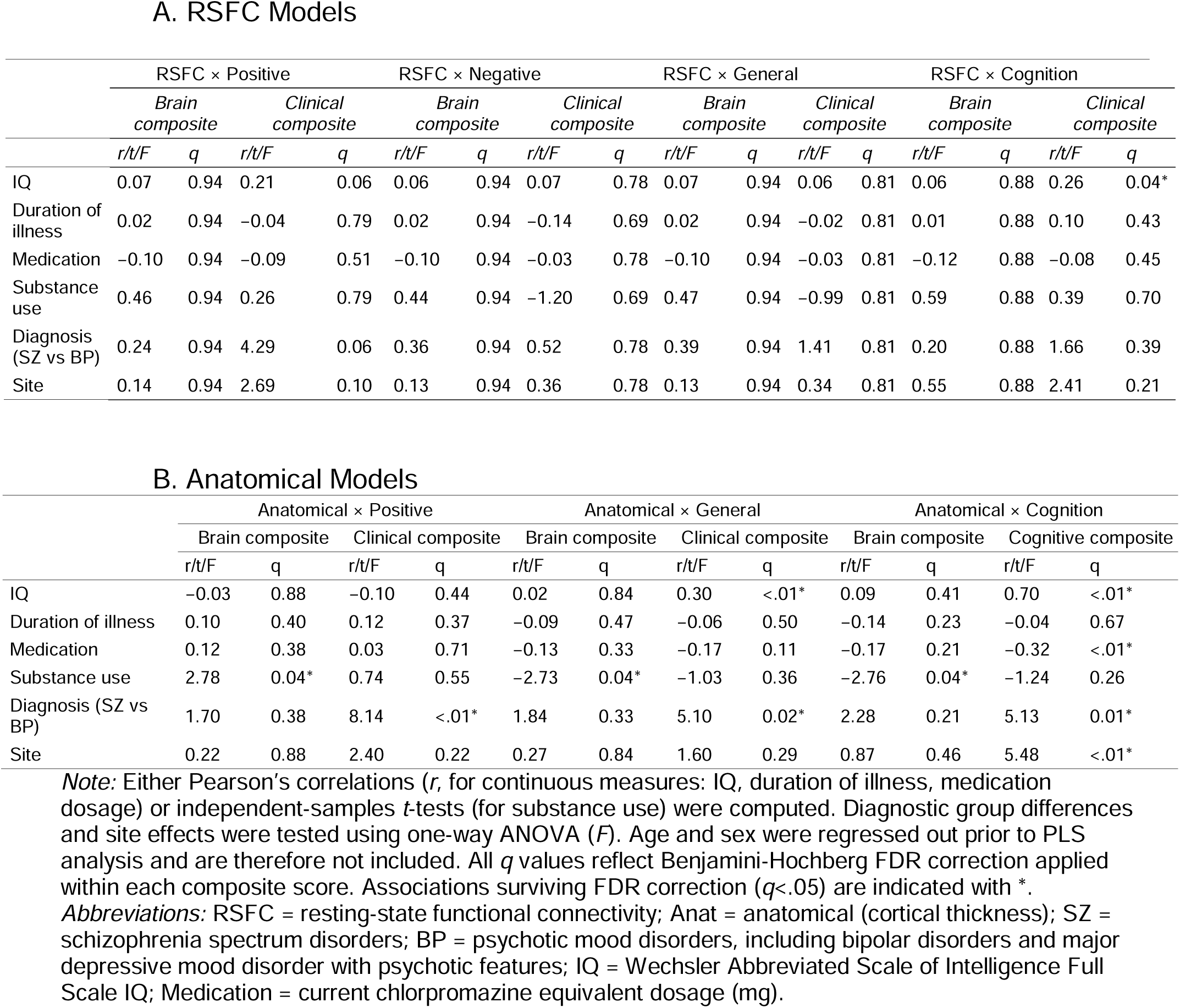
Associations between PLS composite scores and potential confounds across all significant latent components.

### Brain Imaging Measures

#### Resting-State Functional Connectivity

Resting-state fMRI data were preprocessed following standard HCP pipelines, including visual inspection for excessive motion and ICA-AROMA denoising. Full preprocessing details are described in Wang et al., 2025 [22]. Global signal regression was not applied, as the global signal has been shown to correlate with clinical and behavioral measures in psychosis such that its removal may eliminate disease-related variance [29, 30]. Our prior study using resting-state fMRI methods in this cohort found that secondary analyses with global signal regression did not alter interpretation of results [22].

RSFC was computed as Pearson correlations among mean time series of 200 cortical parcels from the Schaefer atlas, corresponding to a subdivision of 17 functional networks [31], and 16 subcortical parcels from the Tian atlas [32], yielding a 216 × 216 connectivity matrix per participant. The 23,220 unique pairwise RSFC strength values served as brain features for PLSc analyses between RSFC and symptoms or cognition.

#### Cortical Thickness and Subcortical Volume

T1-weighted structural images were acquired using an MPRAGE sequence on a 3T Siemens Prisma scanner (TR=2400ms, TE=2.22ms, TI=1000ms, flip angle=8°, 0.8mm isotropic resolution). Structural data were processed using HCP pipelines implemented in QuNex [33]. Cortical thickness was extracted from 68 regions (34 per hemisphere) using the Desikan-Killiany atlas (aparc), and subcortical volumes were extracted from 14 structures (7 per hemisphere) using automated segmentation (aseg). Cortical thickness was selected over surface area as the primary cortical metric given its greater sensitivity to psychosis-related alterations [34–36]. The 82 regional anatomical measures served as brain features for PLSc analyses between anatomical features and symptoms or cognition.

### Clinical and Cognitive Measures

#### Clinical Measures

Four dimensions of psychopathology were assessed. Positive symptoms (7 items), negative symptoms (7 items), and general psychopathology (16 items) were measured using the Positive and Negative Syndrome Scale (PANSS) [37]. Manic symptoms (11 items) were measured using the Young Mania Rating Scale (YMRS) [38]. Each symptom dimension served as a separate behavioral feature set for PLSc analyses, such that item-level scores within each dimension were paired with brain features to derive dimension-specific brain signatures.

#### Cognitive Measures

Cognition was assessed using the NIH Toolbox Cognition Battery [39] and the Wechsler Abbreviated Scale of Intelligence Second Edition (WASI-II) [40]. Age-adjusted standard scores from 11 individual tasks were z-transformed across the sample and averaged within seven cognitive domains based on established frameworks in the schizophrenia cognition literature [41]: general cognitive ability (WASI Full Scale IQ), working memory (Picture Sequence Memory, List Sorting), processing speed (Pattern Comparison), sensorimotor functioning (Grip Strength, 9-Hole Pegboard Dexterity), language (Picture Vocabulary, WASI Vocabulary, Oral Reading Recognition), visuospatial ability (WASI Matrix Reasoning), and executive functioning (Dimensional Change Card Sort, Flanker). The seven domain composite scores served as the cognitive dimension for PLSc analyses.

### Dimension-Specific Brain Signatures

We employed Partial Least Squares correlation (PLSc) using pyls in Python [42] to identify multivariate brain signatures maximally associated with each clinical and cognitive dimension. PLSc decomposes brain and behavioral data matrices into latent components (LCs) that capture maximum covariance between the two domains [43]. Both RSFC strength and anatomical features were harmonized for site effects using ComBat-GAM (NeuroHarmonize) [44]. Prior to analysis, sex, age, and quadratic age were regressed from the brain features using ordinary least squares regression.

We examined both functional and structural modalities as a proof-of-concept approach, given that it remains unclear which brain features best capture dimensional psychopathology or yield informative receptor associations. Ten separate PLSc models were run, pairing each of five dimensions (positive symptoms, negative symptoms, general psychopathology, mania, and cognition) with each of two brain modalities (RSFC and anatomical), yielding 5 dimension × 2 modality pairs. Statistical significance of each LC was assessed via permutation testing (10,000 permutations). Bootstrap resampling (5,000 iterations) was used to estimate confidence intervals for brain and clinical/cognitive loadings; loadings with 95% confidence intervals not crossing zero were considered significant. Full methodological details are provided in our previous work [21, 22].

### Neurotransmitter Receptor Annotation

To identify neurotransmitter receptor systems with spatial distribution that are correlated with the dimension-specific brain signatures, we used the *neuromaps* toolbox [45] to correlate each significant PLSc-derived brain signature with normative maps of neurotransmitter receptor and transporter densities [19]. Forty-one receptor and transporter density maps were derived from positron emission tomography (PET) data aggregated across more than 1,600 healthy adult individuals [23]. We averaged PET maps for the same receptors/transporters to increase the sample size of each receptor/transporter map and reduce noise, after confirming the similarity across maps for the same systems across different tracers (see Supplemental Methods). We examined 21 receptors and transporters spanning nine neurotransmitter systems, including dopamine, norepinephrine, serotonin, acetylcholine, glutamate, GABA, histamine, cannabinoid, and opioid. See Supplementary Methods for details on PET maps. For each significant LC, we computed Spearman correlations between the PLSc-derived brain signature and each receptor/transporter density map. Statistical significance was assessed using spin permutation testing (10,000 permutations) [46], which generates null distributions by applying random rotations to spherical projections of the cortical surface, thereby preserving spatial autocorrelation [46–48]. Results were corrected for multiple comparisons using FDR correction across all receptors.

### Sensitivity Analyses

To evaluate whether PLSc-derived brain–behavior associations were confounded by demographic, clinical, or methodological factors, we conducted sensitivity analyses for all significant LCs. For each LC, we computed Pearson’s correlations between PLSc composite scores and potential confounds, including IQ, duration of illness, antipsychotic medication (chlorpromazine equivalents), substance use, diagnosis (affective vs non-affective psychosis), and acquisition site. Statistical significance was assessed using FDR correction across all confounds.

## RESULTS

### Sample Characteristics

The primary symptom analysis included 121 patients from the HCP-EP cohort (mean age 22.7±3.7 years; 61.2% male). Participants were diagnosed with schizophrenia spectrum disorders (65.3%), schizoaffective disorder (11.6%), or psychotic mood disorders (23.1%), all within three years of illness onset. The cognition analysis included a subset of 111 patients with available cognition data. The cognition subsample did not differ from the full sample on any demographic, clinical, or medication variable (all p>.05). Complete demographic, clinical, and cognitive characteristics are presented in **Table 1**.

### Dimension-Specific Brain Signatures

#### RSFC Strength Signatures

PLS correlation identified significant latent components (LCs) linking whole-brain RSFC to four of the five clinical dimensions tested. Specifically, significant LCs were identified for positive symptoms (LC_pos-RSFC_; 54.8% covariance explained, p=.010), negative symptoms (LC_neg-RSFC_; 49.7%, p=.002), general psychopathology (LC_gps-RSFC_; 44.2%, p=.003), and cognition (LC_cog-RSFC_; 54.2%, p=.002). The mania model did not yield a significant LC (p>.05). In all significant models, brain and clinical composite scores were positively correlated (LC_pos-RSFC_: r=.28, p=.002, **Figure 1A**; LC_neg-RSFC_: r=.31, p<.001, **Figure 1D**; LC_gps-RSFC_: r=.31, p<.001, **Figure 1G**; LC_cog-RSFC_: r=.34, p<.001, **Figure 1J**), reflecting a pattern where greater RSFC strength covaried with greater impairment across symptom and cognitive domains. Composite scores were unrelated to all tested confounds after FDR correction (q>.05).

**Figure 1.**
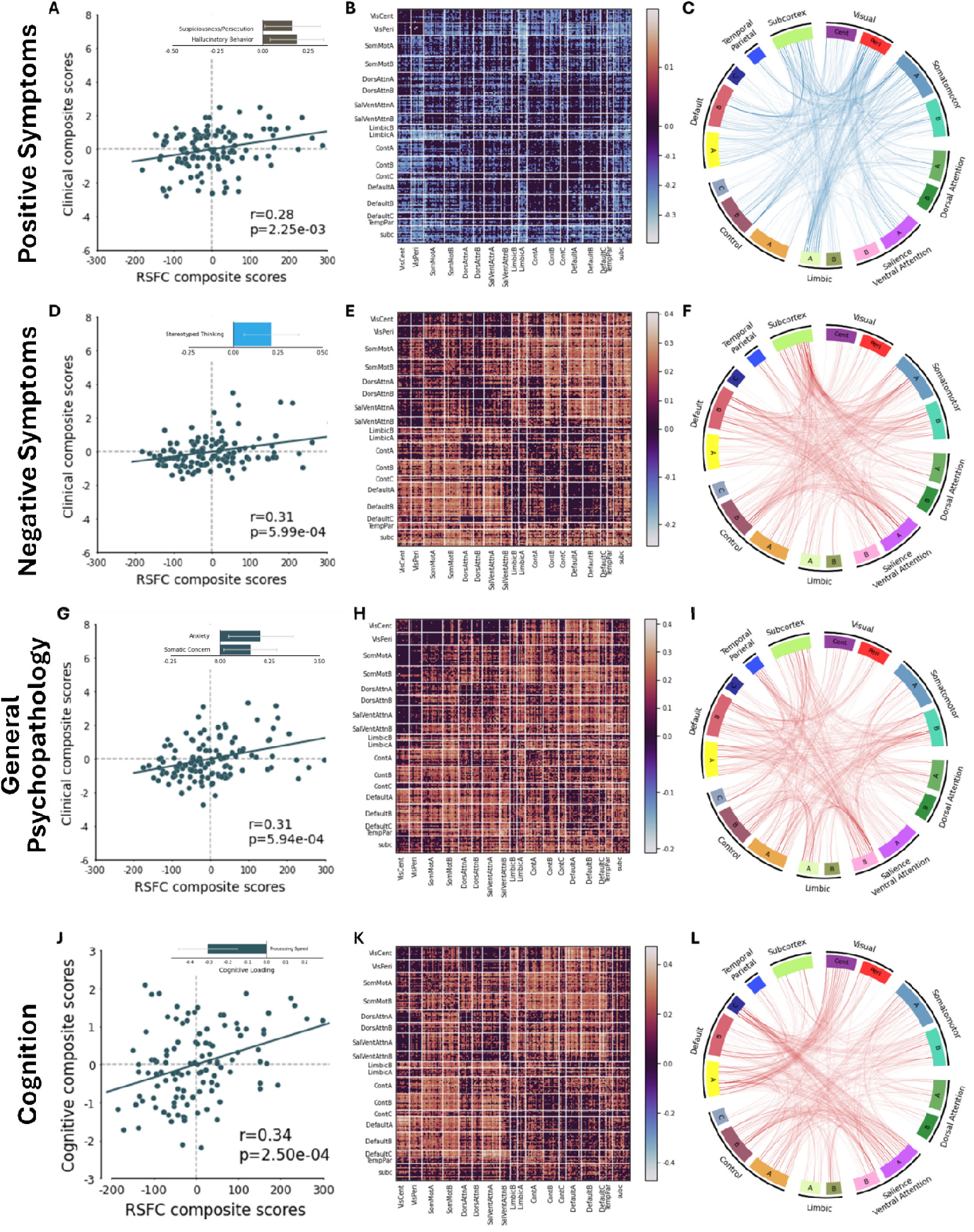
PLSc-derived latent components linking whole-brain RSFC strength to severity of positive symptoms, negative symptoms, general psychopathology, and cognition impairment. Each row presents a significant LC, with scatterplots (left) showing the correlation between RSFC and clinical/cognitive composite scores with inset bar plots displaying stable clinical or cognitive loadings, heatmap matrices (middle) showing unthresholded network loading patterns, and circos plots (right) showing stable network loadings. Only loadings with bootstrap confidence intervals not crossing zero are displayed. In all scatter plots, regression lines are plotted. In all network matrices, red (or blue) color indicates that greater RSFC strength is positively (or negatively) associated with the LC composite score. (**A**) Scatter plot of RSFC and clinical composite scores for the positive symptom model (r=.28, p=.002). Inset bar plot shows bootstrap ratios for stable clinical loadings (suspiciousness and hallucinatory behavior). (**B**) Unthresholded correlations between participants’ RSFC data and their RSFC composite scores for LC_pos-RSFC_ averaged within and between networks. (**C**) Circos plot depicting stable within- and between-network RSFC loadings. Each line represents a stable RSFC loading. (**D**) Scatter plot for the negative symptom model (r=.31, p<.001). Stereotyped thinking was the primary stable clinical loading. (**E**) Unthresholded correlations between participants’ RSFC data and their RSFC composite scores for LC_neg-RSFC_ averaged within and between networks. (**F**) Circos plot of stable RSFC loadings for the negative symptom LC. (**G**) Scatter plot for the general psychopathology model (r=.31, p<.001). Anxiety and somatic concern were the primary stable clinical loadings. (**H**) Unthresholded correlations between participants’ RSFC data and their RSFC composite scores for LC_gps-RSFC_ averaged within and between networks. (**I**) Circos plot of stable RSFC loadings for the general psychopathology LC. (**J**) Scatter plot for the cognition model (r=.34, p<.001). Processing speed negatively loaded onto the clinical composite score of LC_cog-RSFC_. (**K**) Unthresholded correlations between participants’ RSFC data and their RSFC composite scores for LC_cog-RSFC_ averaged within and between networks. (**L**) Circos plot of stable RSFC loadings for the cognition LC.

Bootstrap analyses revealed significant clinical loadings for each LC, identifying which specific symptoms or cognitive domains most reliably drove each brain–behavior relationship. For the positive symptom model, hallucinations (r=.19) and suspiciousness (r=.16) emerged as stable loadings, with the corresponding brain signature reflecting regions where lower RSFC strength covaried with greater positive symptom severity across visual, limbic, and somatomotor networks (**Figure 1A-C**). For negative symptoms, stereotyped thinking (r=.21) emerged as the primary significant loading, with the corresponding brain signature reflecting higher RSFC strength across the subcortical, salience/ventral attention, and control networks (**Figure 1D-F**). The general psychopathology LC was predominantly driven by anxiety (r=.20) and somatic concern (r=.15), with the corresponding brain signature reflecting higher RSFC strength across a widespread set of networks (r<=.40; **Figure 1G-I**). The cognition RSFC model most strongly loaded on lower processing speed (r=-.31), with the corresponding brain signature reflecting higher RSFC strength across the default mode, salience/ventral attention, and central visual networks (**Figure 1J-L**).

#### Anatomical Signatures

Anatomical PLS models identified significant LCs for three of the five dimensions: positive symptoms (LC_pos-anat_; 59.4% covariance explained, p=.01), general psychopathology (LC_gps-anat_; 42.1%, p<.001), and cognition (LC_cog-anat_; 60.2%, p=.014). Neither negative symptoms nor mania yielded significant anatomical LCs (p>.05). In all significant models, brain and clinical composite scores were positively correlated (LC_pos-anat_: r=.29, p=.002, **Figure 2A**; LC_gps-anat_: r=.36, p<.001, **Figure 2C**; LC_cog-anat_: r=.31, p=.001, **Figure 2E**), and were unrelated to confounds (q>.05).

**Figure 2.**
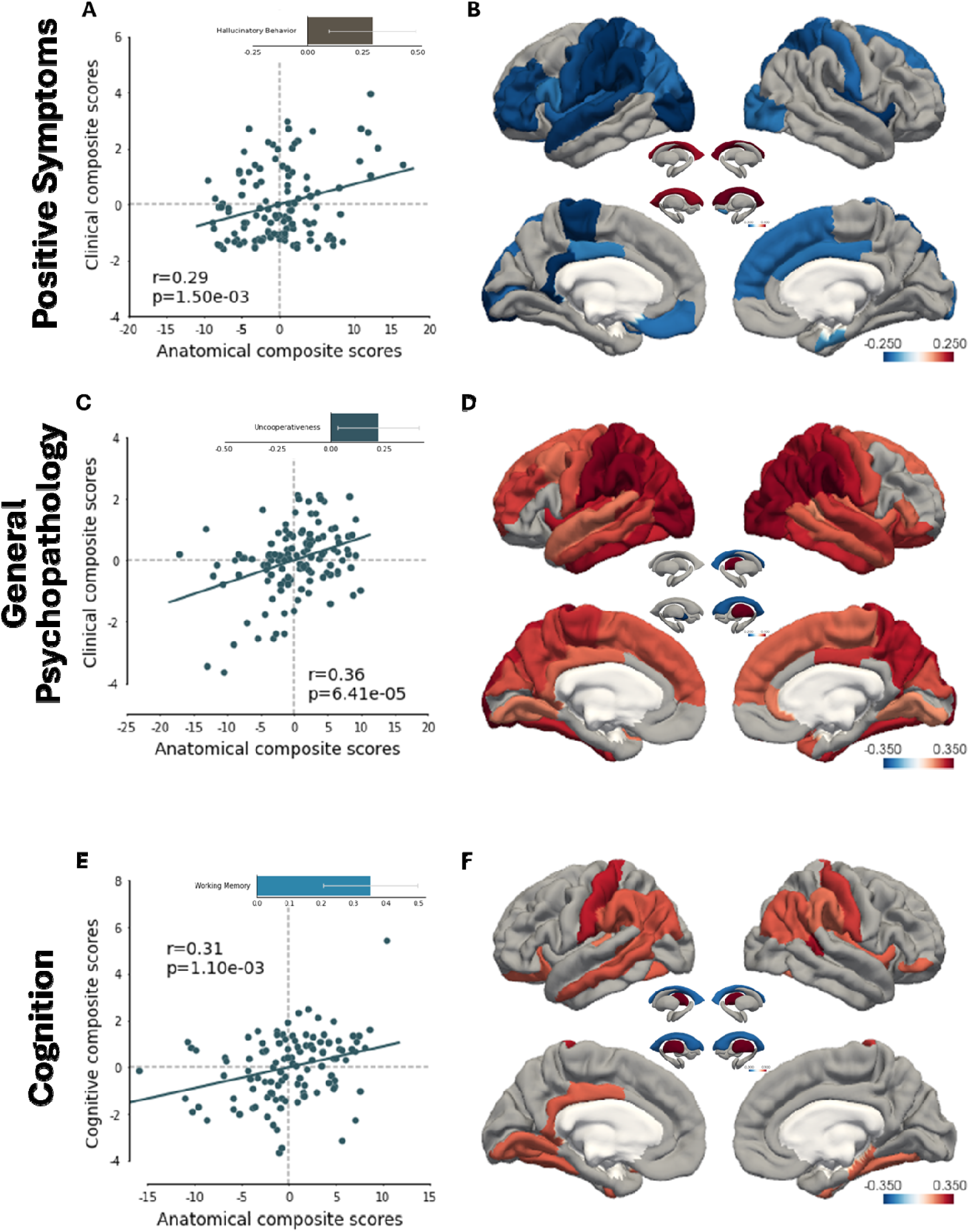
PLSc-derived latent components linking whole-brain anatomical pattern to severity of positive symptoms, general psychopathology, and cognition impairment. Eac row presents a significant anatomical LC, with scatterplots (left) showing the correlation between anatomical and clinical/cognitive composite scores with inset bar plots displaying significant clinical or cognitive loadings, and cortical surface maps with subcortical volume renderings (right) showing regional loading patterns. Warm colors (red) indicate positive loadings (cortical thickening or larger subcortical volume); cool colors (blue) indicate negative loadings (cortical thinning or smaller volume). Only loadings with bootstrap confidence intervals not crossing zero are displayed. (**A**) Scatterplot for LC_pos-anat_ (r=.29, p=.002), with hallucinatory behavior as the stable clinical loading. (**B**) Cortical thinning in bilateral insula, left postcentral gyrus, left isthmus cingulate, and left lateral occipital cortex, with larger bilateral lateral ventricle volumes. (**C**) Scatterplot for LC_gps-anat_ (r=.36, p<.001), with uncooperativeness as the stable clinical loading. (**D**) Widespread cortical thickening in supramarginal, postcentral, inferior/superior parietal, and lateral occipital regions bilaterally, with smaller left pallidum and right lateral ventricle volumes and larger right thalamus volumes. (**E**) Scatterplot for LC_cog-anat_ (r=.31, p=.001), with working memory as the stable cognitive loading. (**F**) Widespread cortical thickening concentrated in posterior association cortex, with negative bilateral lateral ventricle loadings.

Clinical loadings for the anatomical models differed from their RSFC counterparts. LC_pos-anat_ was primarily driven by hallucinations (r=.29), with the brain pattern reflecting widespread cortical thinning in bilateral insula, left postcentral gyrus, left isthmus cingulate, and left lateral occipital cortex (-.27<=r<=-.15, **Figure 2B**). Additionally, larger bilateral lateral ventricles positively loaded onto the anatomical composite scores (- .20<=r<=.32, **Figure 2B**). LC_gps-anat_ was stably associated with the clinical loading of uncooperativeness (r=.22), with widespread cortical thickening across supramarginal gyrus, postcentral gyrus, inferior/superior parietal, and lateral occipital regions bilaterally (.15<=r<=.34, **Figure 2D**). Subcortically, smaller volumes of left pallidum and right lateral ventricles, as well as larger volumes of right thalamus, stably loaded onto the anatomical signature (-.24<=r<=.24, **Figure 2D**). LC_cog-anat_ was positively associated with the loading of working memory (r=.35), and the anatomical signature showed a widespread cortical thickening pattern, similar to the general psychopathology pattern, concentrated in posterior association cortex, while bilateral lateral ventricles showed negative loadings, meaning larger ventricles relate to worse cognitive performance (-.24<=r<=.37, **Figure 2F**).

### Neurotransmitter Receptor Annotation

Spatial correlations between significant brain signatures and 21 neurotransmitter receptor/transporter maps from *neuromaps* revealed dimension-specific molecular associations for RSFC, but not anatomical signatures (**Figure 3**).

**Figure 3.**
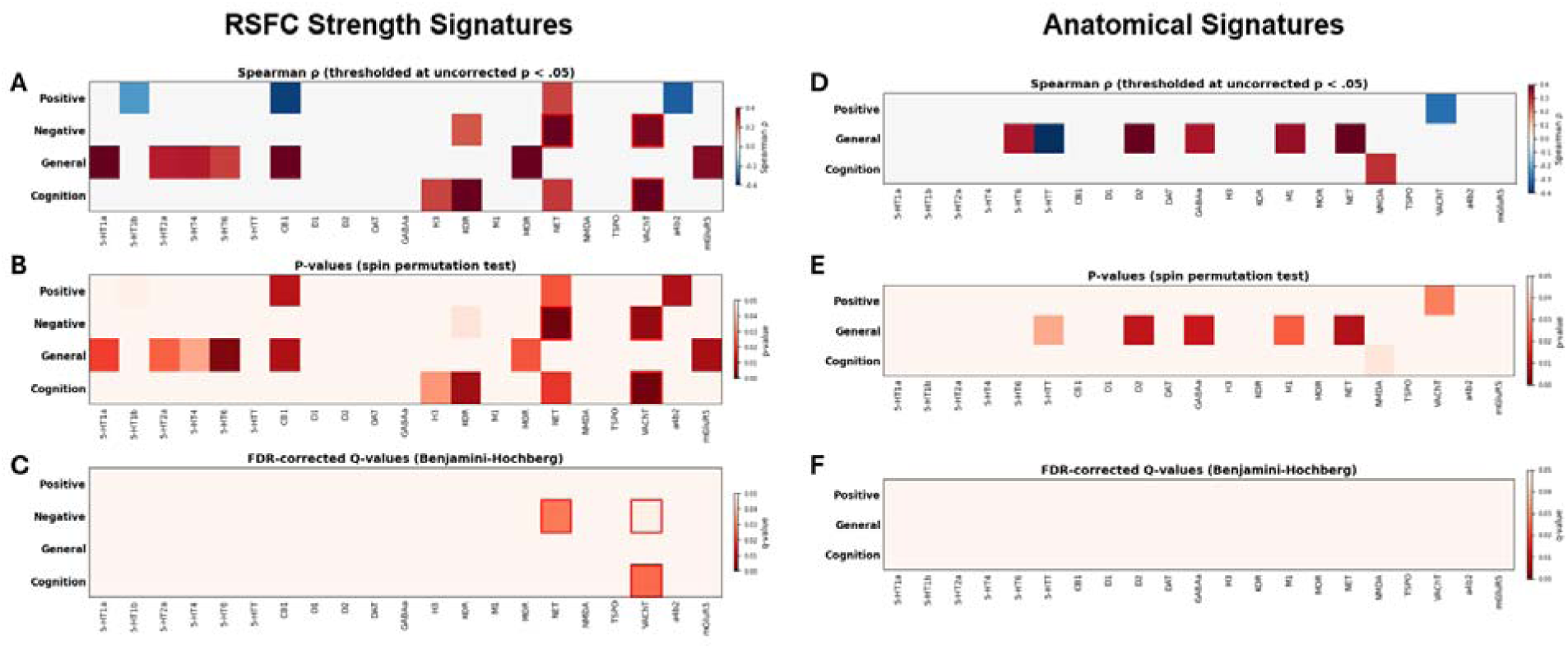
Neurotransmitter Receptor Annotations of RSFC and Anatomical Brain Signatures for Psychopathology Dimensions. Spatial correlations between significant brain signatures and 21 neurotransmitter receptor/transporter density maps are displayed for RSFC (left) and anatomical (right) modalities across symptom and cognition dimensions. (**A**) Spearman correlations (ρ) between RSFC signatures and receptor/transporter maps, thresholded at uncorrected p<.05. (**B**) Corresponding p-values from spin permutation tests (10,000 permutations) for RSFC signatures. (**C**) FDR-corrected q-values (Benjamini-Hochberg) for RSFC signatures. The negative symptom RSFC signature was significantly associated with the norepinephrine transporter (NET; ρ=.40, q=.030) and vesicular acetylcholine transporter (VAChT; ρ=.38, q=.048), while the cognition RSFC signature was associated with VAChT (ρ=.48, q=.025). (**D**) Spearman correlations between anatomical signatures and receptor/transporter maps, thresholded at uncorrected p<.05. (**E**) Corresponding spin permutation test p-values for anatomical signatures. (**F**) FDR-corrected q-values for anatomical signatures; no anatomical associations survived correction (q>.05). Rows represent symptom/cognition dimensions; columns represent receptor/transporter systems. Darker colors indicate stronger correlations (A, D) or smaller p/q-values (B, C, E, F).

#### Receptor Annotation of RSFC Strength Signatures

Among the four RSFC LCs that reached statistical significance, two showed spatial correlations with receptor/transporter maps that survived that FDR correction (q<.05; **Figure 3A-C**). The RSFC signature associated with negative symptoms (LC_neg-RSFC_) was positively spatially correlated with the norepinephrine transporter (NET; ρ=.40, p_spin_=.001, q=.030) and vesicular acetylcholine transporter density maps (VAChT; ρ=.38, p_spin_=.005, q=.048). The cognition RSFC signature (LC_cog-RSFC_) was similarly spatially correlated with VAChT (ρ=.48, p_spin_=.001, q=.025). Notably, the convergence of VAChT across both negative symptoms and cognitive impairment suggests a shared cholinergic substrate linking these two domains.

At an uncorrected threshold (p_spin_<.05), additional associations were observed. Full receptor annotation results for all dimensions are presented in **Figure 3** and Supplementary Table S2-3.

#### Receptor Annotation of Anatomical Signatures

None of the three significant anatomical LCs (positive symptoms, general psychopathology, cognition) displayed significant spatial alignment with any receptor or transporter map after FDR correction (*q*>.05). While some uncorrected associations were observed (*p*_spin_<.05; **Figure 3D-F**), these did not survive multiple comparison correction. This divergence between modalities suggests that RSFC, which putatively reflects dynamic neural activity, may be more suitable than static brain morphology for detecting dimension-specific neurotransmitter associations in early psychosis [49–51].

### Sensitivity Analyses

To assess whether the PLS-derived composite scores were influenced by potential confounds, we examined the associations between each significant LC’s brain and clinical (or cognitive) composite scores and potential confounds, including IQ, duration of illness, antipsychotic medication, substance use, diagnosis, and site. Across all three symptom × RSFC models (LC_pos-RSFC_, LC_neg-RSFC_, LC_gps-RSFC_), neither the RSFC nor clinical composite scores were significantly associated with any confound after FDR correction (all q>.05). For the cognition × RSFC model (LC_cog-RSFC_), the cognitive composite score was associated with IQ (r=.26, q=.04), while the brain composite score was not associated with any confound (all q>.05).

For the symptom × anatomical models, the brain composite scores for both the positive and general psychopathology LCs were associated with substance use (t=2.78, q=.04), and the general psychopathology clinical composite score was correlated with IQ (r=.30, q<.01) and differed by diagnostic group (F=5.10, q=.02). The positive symptom clinical composite score also differed by diagnostic category of affective vs non affective psychosis (F=8.14, q<.01). For the cognition × anatomical model, the cognitive composite score was significantly associated with IQ (r=.70, q<.01), antipsychotic medication dosage (r=−.32, q<.01), diagnostic group (F=5.13, q=.01), and site (F=5.48, q<.01), while the brain composite score was associated only with substance use (q=.04).

## DISCUSSION

This study identified dimension-specific whole-brain signatures associated with positive symptoms, negative symptoms, general psychopathology, and cognition in a transdiagnostic early psychosis cohort, and characterized their spatial correspondence with neurotransmitter receptor and transporter distributions. In showing that distinct symptom dimensions map onto distinct brain circuits with distinct receptor correlates, our findings support a dimensional rather than categorical view of psychosis neurobiology. Receptor annotation revealed that the negative symptom RSFC signature was significantly associated with norepinephrine transporter (NET) and vesicular acetylcholine transporter (VAChT) distributions, while the cognition RSFC signature was spatially correlated with VAChT distribution. Critically, the RSFC brain signatures were unrelated to antipsychotic medication and substance use, strengthening confidence that receptor associations reflect intrinsic brain-symptom relationships rather than confounds. Anatomical signatures did not yield significant receptor associations, and were more susceptible to confounding by clinical variables. These findings are consistent with the implication of cholinergic and noradrenergic systems in the neurobiology of negative symptoms and cognitive impairment in early psychosis.

The identified brain signatures demonstrated face validity with established neuroscience of each symptom domain. The negative symptom RSFC pattern, characterized by elevated functional connectivity in subcortical, salience/ventral attention, and frontoparietal control networks, aligns with models implicating cortico-striatal-thalamic circuits in motivational deficits and goal-directed behavior [52]. The cognition RSFC pattern showed elevated functional connectivity in default mode and salience/ventral attention networks, consistent with evidence that alterations in network interactions, and failure to suppress default mode activity during task performance impairs cognitive efficiency [53]. The positive symptom pattern, marked by reduced functional connectivity in visual, limbic, and somatomotor networks, aligns with findings implicating sensory network dysconnectivity in hallucinations [54]. Anatomically, the positive symptom signature—cortical thinning in insula and cingulate with ventricular enlargement—is consistent with some of the canonical morphometric findings in psychosis [34]. These patterns provide biologically plausible substrates for subsequent neurotransmitter annotation.

VAChT emerged as the only receptor/transporter significantly associated with both negative symptom and cognition RSFC signatures, suggesting that cholinergic terminal density represents a shared neurobiological substrate. This convergence aligns with the well-documented clinical overlap between negative symptoms and cognitive deficits in schizophrenia, both of which remain treatment-resistant and predictive of functional outcomes [11, 55, 56]. VAChT localizes to presynaptic terminals of cholinergic projection neurons originating in the basal forebrain and brainstem [57, 58], and its spatial distribution may reflect the integrity of cholinergic inputs to cortical and subcortical targets. Our findings are consistent with prior PET studies demonstrating that VAChT binding in dorsolateral prefrontal cortex was inversely correlated with working memory performance in patients with schizophrenia [59], establishing a direct link between presynaptic cholinergic integrity and cognition. Although M1 muscarinic receptor maps did not show significant associations, this null result should be interpreted cautiously given methodological limitations of early-generation M1 PET tracers [60] and the map quality variation of early small sample studies in *neuromaps*.

Importantly, clinical research strongly supports muscarinic involvement in psychosis [61, 62]. For example, Cobenfy (xanomeline-trospium), a muscarinic M1/M4 agonist, recently received FDA approval for schizophrenia treatment based on demonstrated efficacy for positive, negative, and cognitive symptoms [17, 63]. Our VAChT findings and the clinical efficacy of muscarinic agonists together implicate cholinergic signaling as a substrate for negative symptoms and cognitive deficits warranting further mechanistic investigation.

The association between the negative symptom RSFC signature and norepinephrine transporter NET density maps represents, to our knowledge, the first neuroimaging evidence linking noradrenergic transporter distribution to negative symptoms in psychosis. To our knowledge, no prior PET studies have examined NET in schizophrenia due to lack of reliable PET radioligands [50, 64], making the present finding hypothesis-generating and warranting replication with direct molecular imaging. Noradrenergic signaling via locus coeruleus projections modulates arousal, attention, and prefrontal network function [65–68], domains consistently impaired in schizophrenia and particularly relevant to negative symptoms. NET serves as the primary transporter for both norepinephrine and dopamine in the prefrontal cortex, where dopamine transporter (DAT) expression is sparse [69, 70]. Consequently, NET dysfunction could affect prefrontal catecholamine clearance and contribute to the motivational and cognitive deficits characteristic of negative symptoms [71]. Taken together, our findings provide a neurobiological rationale for systematic evaluation of NET-targeting compounds for negative symptoms in psychosis.

Several null findings merit consideration. Dopamine receptor distributions (D1, D2) did not significantly correlate with any brain signature, despite dopamine’s central role in psychosis pathophysiology. This apparent discrepancy likely reflects methodological rather than biological considerations. Dopamine dysfunction in psychosis is primarily presynaptic, involving elevated synthesis capacity and release rather than altered postsynaptic receptor density [72–74]. The *neuromaps* receptor atlas captures postsynaptic receptor expression, not presynaptic dynamics (see Supplemental Methods). Additionally, the D1 and D2 receptors exhibit highest density in the striatum with relatively less cortical spatial variance, limiting detection sensitivity in whole-brain spatial correlation analyses [75]. Glutamate and GABA receptor distributions similarly showed no significant associations, despite their importance in psychosis neurobiology [76, 77]. Prior work using MEG found that ionotropic glutamate and GABA receptors were among the most dominant for oscillatory dynamics [19], suggesting that these systems may be better captured by electrophysiological measures of neural activity than by RSFC. Alternatively, glutamatergic and GABAergic dysfunction in psychosis may be more regionally localized, rather than following whole-brain gradients amenable to spatial correlation [78, 79]. Similarly, α4β2 nicotinic receptors showed no significant associations, despite longstanding interest in nicotinic signaling in schizophrenia, stemming from elevated smoking rates [80, 81] and evidence of α7 nicotinic receptor deficits in patients with schizophrenia [82, 83].

However, the α4β2 map in *neuromaps* was derived from a single study, limiting power to detect associations. Hence this null finding should be interpreted cautiously, pending availability of larger normative samples. Furthermore, the mania dimension yielded no significant PLSc brain signature in either modality, precluding receptor annotation. This null finding likely reflects sample composition and symptom characteristics, where the HCP-EP cohort is predominantly schizophrenia spectrum (65.3%), and most participants were not experiencing symptoms of elevated mood at scan, producing floor effects on the mania symptom severity.

Anatomical signatures showed different clinical loadings than their RSFC counterparts and were more susceptible to confounding by substance use, medication, diagnosis, and site. This pattern suggests that static brain morphology may be more vulnerable to chronic illness effects and less sensitive to dimension-specific neurotransmitter associations than dynamic functional connectivity measures. The absence of any significant receptor correlations for anatomical signatures further supports RSFC as the more suitable modality for this PLSc-mediated analytic approach in early psychosis.

Several limitations should be acknowledged. First, PET-derived receptor maps in the *neuromaps* atlas have variable sample sizes across systems, potentially affecting statistical power to detect associations for less well-characterized receptors. Second, our moderate sample size necessitates replication in larger cohorts before clinical translation. Third, spatial correlation provides annotation rather than direct evidence of mechanistic involvement, though this approach represents the current methodological standard for linking macroscale brain patterns to molecular architecture. Finally, this cross-sectional design cannot address whether receptor associations reflect trait vulnerability, state-dependent changes, or illness progression. Future studies should replicate these findings in independent early psychosis samples with careful attention to medication status. Application of this paradigm to clinical high-risk populations could also help determine whether cholinergic and noradrenergic associations emerge prior to psychosis onset, potentially informing early intervention targets. Because the framework is modality-generalizable, it extends naturally beyond RSFC and structural MRI to diffusion and task MRI, as well as electrophysiological modalities such as electroencephalogram (EEG) or magnetoencephalography (MEG), enabling systematic investigation of how molecular architecture relates to different aspects of neurodynamics, given evidence that different receptor systems preferentially influence different neural signal types.

## CONCLUSIONS

These findings implicate cholinergic (VAChT) and noradrenergic (NET) systems as molecular targets with psychopharmacological potential for treating negative symptoms and cognitive impairment. By bridging dimensional psychopathology with molecular neuroanatomy, this approach provides data-driven, complementary evidence to studies of direct molecular mechanisms, investigating novel pharmacological interventions.

## DATA AVAILABILITY

The HCP-EP dataset is available at https://db.humanconnectome.org/. To facilitate reproducibility and rigor, all analysis code is publicly available on GitHub: https://github.com/haleyrwang/CCNL_Symptoms_Neurotransmitters. A pre-print of this manuscript prior to peer review is hosted on bioRxiv.

## ACKNOWLEDGEMENT

Research using Human Connectome Project for Early Psychosis (HCP-EP) data reported in this publication was supported by the National Institute of Mental Health of the National Institutes of Health under Award Number U01MH109977. This work used computational and storage services associated with the Hoffman2 Shared Cluster provided by UCLA Institute for Digital Research and Education’s Research Technology Group.

## AUTHOR CONTRIBUTIONS

HRW had full access to all of the data in the study and takes responsibility for the integrity of the data and the accuracy of the data analysis.

Study concept and design: HRW, CHS, ZQL, CEB, KHK. Acquisition, analysis, or interpretation of data: HRW, CHS, ZQL, RAM, RB, CMA, HF, LQU, CEB, KHK. Drafting of the manuscript: HRW, CHS, KHK. Critical revision of the manuscript for important intellectual content: HRW, CHS, ZQL, LQU, CEB, BM, KHK. Statistical analysis: HRW, CHS, ZQL, KHK. Obtained funding: KHK. Administrative, technical, or material support: CHS, ZQL, RB, LQU, CEB, BM, KHK. Study supervision: HRW, KHK.

## FUNDING

KHK and CEB reported grants from the National Institute of Mental Health (NIMH). HRW reported support from a COGDOP scholarship from the American Psychological Foundation. LQU is supported by National Institute of Child Health and Human Development (NICHD). CMA is supported by National Institute on Drug Abuse (NIDA).

## COMPETING INTERESTS

The authors declare no competing interests.

## Notes

### Competing Interest Statement

The authors have declared no competing interest.

