## Supplemental Data 1 for "Symptom Dimension-Specific Neurotransmitter Correlates of Psychopathology and Cognition in Early Psychosis"

***Supplemental Information***

**Supplemental Methods**

**Methods S1.** Participant details

**Methods S2.** Categorization and dosage of medication

**Methods S3.** Substance use

**Methods S4.** MRI data acquisition

**Methods S5.** Resting-state imaging data analysis

**Methods S6.** RSFC Parcellation

**Methods S7.** Structural imaging data analysis

**Methods S8.** Partial least squares analysis

**Methods S9.** Neurotransmitter receptor and transporter density maps

**Methods S10.** Averaging and cross-tracer similarity of PET maps

**Supplemental Results**

**Table S1.** Neurotransmitter receptors and transporters used in the *neuromaps* annotation analysis

**Table S2.** Full receptor and transporter annotation results across all dimensions in the RSFC model

**Table S3**. Full receptor and transporter annotation results across all dimensions in the anatomical model

**Supplemental References**

This supplementary material has been provided by the authors to give readers additional information about their work.

***Methods S1. Participant details***

This investigation utilized data from the Human Connectome Project for Early Psychosis (HCP-EP, release 1.1), comprising 182 participants (124 individuals with psychosis and 58 demographically matched unaffected controls) between ages 16-35 at enrollment. The clinical sample included participants meeting diagnostic criteria according to the Diagnostic and Statistical Manual of Mental Disorders (DSM-5) for either primary psychotic disorders (schizophrenia, schizophreniform, schizoaffective, psychosis NOS, delusional disorder, or brief psychotic disorder) or mood disorders with psychotic features (major depression with psychosis or bipolar disorder with psychosis). A key inclusion criterion was psychosis onset within three years prior to study participation. Exclusion criteria encompassed intellectual disability (IQ below 70), history of significant neurological conditions or traumatic brain injury, and contraindications for MRI assessment. Three participants were excluded from neuroimaging analyses due to absent T1-weighted structural scans, and an additional three were excluded from clinical analyses owing to incomplete psychiatric symptom assessments. All assessments were conducted by the HCP research team following protocols detailed in the HCP-EP_Release_1.1_Manual. Symptom evaluation in the psychosis group was performed using the Positive and Negative Syndrome Scale (PANSS) and Young Mania Rating Scale (YMRS).

***Methods S2. Categorization and dosage of medication***

Antipsychotic medication dosages were standardized by conversion to chlorpromazine equivalent values. We examined potential relationships between these standardized medication loads and Partial Least Square correlation (PLSc) composite scores using Pearson correlation analyses (Table 2). The study did not account for other pharmacological treatments such as mood stabilizers, antidepressants, anxiolytics/sedatives/hypnotics, or stimulants, as comprehensive medication information was not available in the public datasets utilized.

***Methods S3. Substance use***

HCP-EP participants underwent substance use screening on the day of their scanning session. Individuals with substance use disorders included in the study presented either mild symptoms or were in remission. For our analyses, substance use was specifically defined as cannabis consumption within the preceding 30 days, a focus selected due to the elevated prevalence of cannabis use among individuals with psychotic disorders. Cannabis use status was categorized dichotomously as either present or absent, without further quantification of usage frequency. We evaluated potential associations between cannabis use status and PLSc composite scores using independent two-sample t-tests (Table 2).

***Methods S4. MRI data acquisition***

Imaging data of the HCP-EP study were collected on three Siemens MAGNETOM Prisma 3T scanners across 4 sites. Scanning sessions included a collection of T1-weighted (multi-echo 3D MPRAGE, TR/TE/TI 2400/2.22/1000ms, 8° flip angle, FOV 256mm, 0.8mm isotropic resolution, 2x GRAPPA acceleration, 208 sagittal slices) and T2-weighted (3D variable-flip-angle turbo-spin-echo (TSE) sequence SPACE, TR/TE 3200/563ms, FOV 256mm, 0.8mm isotropic, 2x GRAPPA, turbo factor 314ms, 208 sagittal slices) structural scans. Additionally, spin echo field maps were collected 4 times with twice for each AP and PA phase encoding (TR/TE 8000/66ms, FoV 208mm, 90° flip angle, 2mm isotropic resolution). Resting state functional MRI data were acquired using a multiband echo-planar imaging sequence (TR/TE 800/37ms, 52° flip angle, FOV 208mm, 2mm isotropic resolution, multiband acceleration factor 8). Four runs were collected, with two runs using anterior-to-posterior and two runs using posterior-to-anterior phase encoding directions. Each run had an acquisition time of 5:47 minutes.

***Methods S5. Resting-state imaging data analysis***

Raw structural and resting-state functional MRI (rs-fMRI) data in NIfTI format were acquired from HCP-EP study. Preprocessing and quality control for HCP-EP were performed using HCP Pipelines(4) implemented through Quantitative Neuroimaging Environment & Toolbox (QuNex)(5), which is compatible with multi-band and single-band fMRI. The fMRI time series underwent additional processing steps including bandpass filtering; motion scrubbing for frames that exceeded either framewise displacement or signal change thresholds; and spatial smoothing. Global signal regression (GSR) was not performed for the main analyses. Three patients were excluded due to missing T1w images. Any scans where more than 50% of frames were flagged for motion were removed from analysis. Additionally, ComBat-GAM (i.e., NeuroHarmonize) was employed to harmonize RSFC data to adjust for scanner difference in the sample.

***Methods S6. RSFC Parcellation***

Brain parcellation methodology. (A). Cortical regions based on Schaefer's 200-parcel atlas. These parcels are organized into 17 resting-state networks [4]. Note that the 200 cortical parcellations are not symmetric between hemispheres. (B). Subcortical structures comprising 16 regions [5]. These combined parcellations yielded 216 × 216 RSFC matrices for each participant. Due to the symmetrical nature of RSFC matrices, analyses utilized only the upper triangular portions (though complete matrices are displayed for visualization purposes).

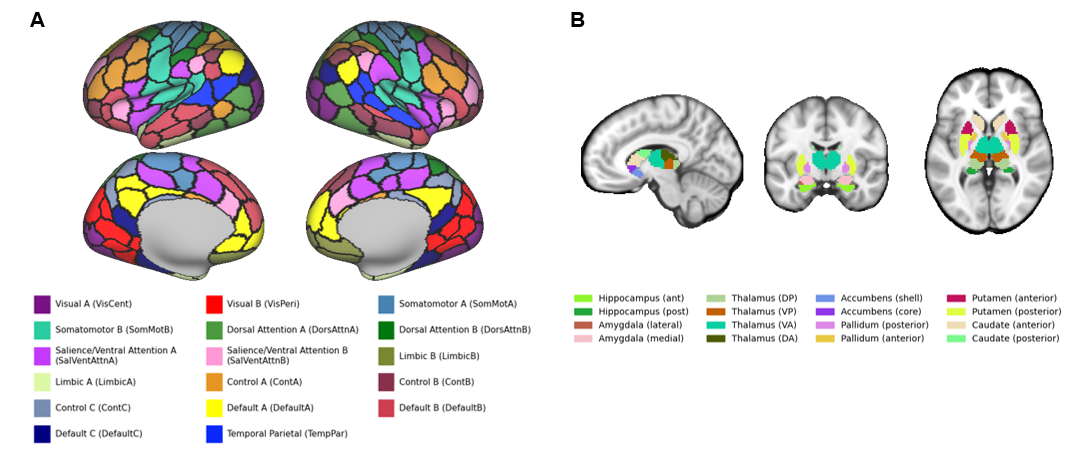

***Methods S7. Structural imaging data analysis***

Anatomical brain features were derived from T1-weighted structural images processed through the HCP structural pipeline implemented in QuNex. For each participant, cortical thickness was estimated for 68 cortical regions (34 per hemisphere) defined by the Desikan-Killiany atlas, and volumes were estimated for 14 subcortical structures (7 per hemisphere) using FreeSurfer’s automated subcortical segmentation (i.e., aseg). Cortical thickness was chosen as the primary cortical metric, rather than surface area or cortical volume, because it shows the greatest sensitivity to psychosis-related morphometric alterations in large consortium studies. This yielded 82 regional anatomical measures per participant (68 cortical thickness values and 14 subcortical volumes), which served as the brain feature set for the anatomical partial least squares correlation (PLSc) analyses.

Prior to analysis, the 82 anatomical features were harmonized across acquisition sites using ComBat-GAM (NeuroHarmonize) to remove scanner- and site-related variance while preserving biological signal, following the same harmonization procedure applied to the RSFC features. Sex, age, and quadratic age were then regressed from each harmonized anatomical feature using ordinary least squares regression, and the residuals were used in subsequent analyses.

***Methods S8. Partial least squares analysis***

We employed partial least squares (PLS) correlation to identify linear relationships between brain RSFC or anatomical features and symptoms or cognition measures using the Pyls 0.1.7 Python package. This unsupervised approach decomposes the cross-product matrix via singular value decomposition, yielding RSFC/anatomical loadings, clinical loadings, and singular values representing relationship strength. Prior to analysis, we controlled for age, quadratic age, and sex through linear regression and z-transformed the residuals.

Feature contribution was assessed using squared-loadings with z-scores above 2 indicating significant impact. Statistical significance was determined through permutation testing (5,000 repetitions) of singular values, while feature stability was evaluated using bootstrap resampling with 95% confidence intervals. Split-half resampling (10,000 iterations) was conducted to examine loading reproducibility. For comprehensive analytical details, see Wang et al. [3].

***Methods S9. Neurotransmitter receptor and transporter density maps***

Normative neurotransmitter receptor and transporter density maps were obtained through the neuromaps toolbox, which aggregates positron emission tomography (PET) images collected from healthy adult volunteers across multiple independent studies [1, 2]. We examined 21 receptors and transporters spanning nine neurotransmitter systems—dopamine, norepinephrine, serotonin, acetylcholine, glutamate, GABA, histamine, cannabinoid, and opioid—selected on the basis of map availability and prior use in molecular-annotation studies. Each PET map was parcellated into the 200-region Schaefer cortical atlas (17-network solution) to match the resolution of the brain signatures prior to spatial correlation. The full list of maps, tracers, and source studies is provided in Table S1.

***Methods S10. Averaging and cross-tracer similarity of PET maps***

Several receptors and transporters were imaged with more than one tracer or in more than one cohort. For each such system, individual maps were first parcellated and then averaged to produce a single representative density map per receptor/transporter, increasing the effective sample size and reducing tracer- and cohort-specific noise (see Table S1 for combined sample sizes). Prior to averaging, we confirmed that maps of the same receptor/transporter acquired with different tracers were spatially consistent by Pearson’s correlation (all p<.05).

**Supplemental Results**

**Table S1. Neurotransmitter receptors and transporters used in the neuromaps annotation analysis.**

| **System** | **Receptor/**  **Transporter** | **Tracer** | **N (males)** | **Age (years)** | **Reference** | **Combined N** |
| --- | --- | --- | --- | --- | --- | --- |
| Serotonin | 5-HT1a | [cumi101] | 8 (3) | 28.4 +/- 8.8 | Beliveau et al., 2017 | 43 |
|  | 5-HT1a | [way100635] | 35 (18) | 26.3 +/- 5.2 | Savli et al., 2012 |  |
|  | 5-HT1b | [az10419369] | 36 (24) | 27.8 +/- 6.9 | Beliveau et al., 2017 | 124 |
|  | 5-HT1b | [p943] | 65 (49) | 28.7 +/- 7 | Gallezot et al., 2010 |  |
|  | 5-HT1b | [p943] | 23 (15) | 28.7 +/- 7 | Savli et al., 2012 |  |
|  | 5-HT2a | [cimbi36] | 29 (15) | 22.6 +/- 2.7 | Beliveau et al., 2017 | 48 |
|  | 5-HT2a | [altanserin] | 19 (11) | 28.2 +/- 5.7 | Savli et al., 2012 |  |
|  | 5-HT4 | [sb207145] | 59 (41) | 25.9 +/- 5.3 | Beliveau et al., 2017 | 59 |
|  | 5-HT6 | [gsk215083] | 30 (30) | 36.6 +/- 9.04 | Radhakrishnan et al., 2018 | 30 |
|  | 5-HTT* | [dasb] | 100 (29) | 25.1 +/- 5.8 | Beliveau et al., 2017 | 128 |
|  | 5-HTT* | [madam] | 10 (8) | 51-67 | Fazio et al., 2016 |  |
|  | 5-HTT* | [dasb] | 18 (12) | 30.5 +/- 9.5 | Savli et al., 2012 |  |
| Dopamine | D1 | [sch23390] | 13 (6) | 33 +/- 13 | Kaller et al., 2017 | 13 |
|  | D2 | [raclopride] | 7 (7) | 24 +/- 2 | Alakurtti et al., 2015 | 304 |
|  | D2 | [fallypride] | 49 (16) | 18.41 +/- 0.57 | Jaworska et al., 2020 |  |
|  | D2 | [raclopride] | 156 (120) | 27.76 +/- 9.44 | Malen et al., 2022 |  |
|  | D2 | [flb457] | 55 (26) | 32.45 +/- 9.69 | Sandiego et al., 2015 |  |
|  | D2 | [flb457] | 37 (17) | 48.36 +/- 16.93 | Smith et al., 2017 |  |
|  | DAT* | [fpcit] | 174 (109) | 61 +/- 11 | Dukart et al., 2018 | 180 |
|  | DAT* | [fepe2i] | 6 (6) | 31.06 +/- 7.7 | Sasaki et al., 2012 |  |
| Norepi-nephrine | NET* | [mrb] | 77 (50) | 33.4 +/- 9.17 | Ding et al., 2010 | 87 |
|  | NET* | [methylre-boxetine] | 10 (n/a) | 33.3 (mean) | Hesse et al., 2017 |  |
| Histamine | H3 | [gsk189254] | 8 (7) | 31.69 +- 8.95 | Gallezot et al., 2017 | 8 |
| Acetylcholine | VAChT* | [feobv] | 18 (5) | 66.8 +/- 6.8 | Aghourian et al., 2017 | 24 |
|  | VAChT* | [feobv] | 5 (4) | 68.3 +/- 3.1 | Bedard et al., 2019 |  |
|  | VAChT* | [feobv] | 4 (3) | 37 +/- 10.2 | Hansen et al., 2022 |  |
|  | a4b2 | [flubatine] | 30 (20) | 33.50 +/- 10.71 | Hillmer et al., 2016 | 27 |
|  | M1 | [lsn3172176] | 24 (13) | 40.45 +/- 11.71 | Naganawa et al., 2020 | 30 |
| Cannabinoid | CB1 | [fmpepd2] | 22 (11) | 27.5 +/- 8.05 | Laurikainen et al., 2019 | 99 |
|  | CB1 | [omar] | 77 (49) | 30.01 +/- 8.87 | Normandin et al., 2015 |  |
| Opioid | MOR | [carfentanil] | 204 (132) | 32.3 +/- 10.8 | Kantonen et al., 2020 | 28 |
|  | MOR | [carfentanil] | 39 (19) | 39.38 +/- 5.05 | Turtonen et al., 2021 |  |
|  | KOR | [ly2795050] | 28 (19) | 33.5 +/- 11.3 | Vijay et al., 2018 | 243 |
| Glutamate | NMDA | [ge179] | 29 (21) | 40.9 +/- 12.7 | Hansen et al., 2022 | 29 |
|  | mGluR5 | [abp688] | 28 (15) | 33.1 +/- 11.2 | Dubois et al., 2016 | 123 |
|  | mGluR5 | [abp688] | 22 (12) | 67.9 +/- 9.6 | Hansen et al., 2022 |  |
|  | mGluR5 | [abp688] | 73 (25) | 19.9 +/- 3.04 | Smart et al., 2019 |  |
| GABA | GABAa | [flumazenil] | 6 (6) | 43 +/- 4 | Dukart et al., 2018 | 32 |
|  | GABAa | [ro154513] | 10 (6) | 25.40 +/- 3.20 | Lukow et al., 2022 |  |
|  | GABAa | [flumazenil] | 16 (7) | 26.6 +/- 8 | Norgaard et al., 2021 |  |
| TSPO | TSPO | [pbr28] | 6 (2) | 57.8 +/- 8.1 | Lois et al., 2018 | 6 |

***Note:*** *Asterisks indicate transporters.* *Where multiple tracers or cohorts were available for the same receptor/transporter, maps were averaged into a single density map (combined N shown in the right-hand column for the first row of each averaged set). Twenty-one receptors and transporters across nine neurotransmitter systems were included in the annotation analysis. TSPO is a translocator-protein glial marker rather than a neurotransmitter system and is listed for completeness but is not counted among the nine systems.*

**Table S2. Full receptor and transporter annotation results across all dimensions in the RSFC model.**

| **Dimension** | **System** | **Receptor/Transporter** | **ρ** | **p_spin** | **q_FDR** |
| --- | --- | --- | --- | --- | --- |
| Positive  Symptoms | Dopamine | D1 | 0.033 | 0.482 | 0.482 |
|  |  | D2 | 0.055 | 0.332 | 0.405 |
|  |  | DAT | 0.175 | 0.201 | 0.405 |
|  | Norepinephrine | NET | 0.27 | 0.022 | 0.154 |
|  | Serotonin | 5-HT2a | 0.026 | 0.446 | 0.469 |
|  |  | 5-HT4 | 0.043 | 0.399 | 0.441 |
|  |  | 5-HT6 | -0.091 | 0.279 | 0.405 |
|  |  | 5-HT1a | 0.097 | 0.339 | 0.405 |
|  |  | 5-HTT | 0.167 | 0.191 | 0.405 |
|  |  | 5-HT1b | -0.23 | 0.048 | 0.252 |
|  | Acetylcholine | VAChT | 0.066 | 0.347 | 0.405 |
|  |  | M1 | -0.116 | 0.211 | 0.405 |
|  |  | α4β2 | -0.33 | 0.008 | 0.102 |
|  | Glutamate | NMDA | 0.075 | 0.303 | 0.405 |
|  |  | mGluR5 | -0.202 | 0.107 | 0.376 |
|  | GABA | GABA-A | -0.064 | 0.341 | 0.405 |
|  | Histamine | H3 | -0.215 | 0.101 | 0.376 |
|  | Cannabinoid | CB1 | -0.374 | 0.01 | 0.102 |
|  | Opioid | KOR | -0.098 | 0.289 | 0.405 |
|  |  | MOR | -0.202 | 0.14 | 0.405 |
|  | TSPO | TSPO | 0.138 | 0.236 | 0.405 |
| Negative Symptoms | Dopamine | D1 | 0.039 | 0.398 | 0.465 |
|  |  | D2 | 0.123 | 0.176 | 0.41 |
|  |  | DAT | 0.141 | 0.212 | 0.41 |
|  | Norepinephrine | NET* | 0.396* | 0.001* | 0.027* |
|  | Serotonin | 5-HT2a | 0.003 | 0.467 | 0.484 |
|  |  | 5-HT1a | 0.062 | 0.392 | 0.465 |
|  |  | 5-HT6 | 0.068 | 0.286 | 0.41 |
|  |  | 5-HT4 | -0.099 | 0.287 | 0.41 |
|  |  | 5-HT1b | 0.114 | 0.236 | 0.41 |
|  |  | 5-HTT | 0.178 | 0.157 | 0.41 |
|  | Acetylcholine | M1 | -0.126 | 0.192 | 0.41 |
|  |  | α4β2 | 0.207 | 0.079 | 0.331 |
|  |  | VAChT* | 0.377* | 0.005* | 0.048* |
|  | Glutamate | mGluR5 | 0.002 | 0.484 | 0.484 |
|  |  | NMDA | 0.068 | 0.293 | 0.41 |
|  | GABA | GABA-A | -0.113 | 0.244 | 0.41 |
|  | Histamine | H3 | 0.238 | 0.062 | 0.327 |
|  | Cannabinoid | CB1 | -0.037 | 0.432 | 0.478 |
|  | Opioid | MOR | 0.078 | 0.313 | 0.411 |
|  |  | KOR | 0.248 | 0.045 | 0.316 |
|  | TSPO | TSPO | 0.121 | 0.214 | 0.41 |
| General | Dopamine | DAT | 0.101 | 0.345 | 0.381 |
|  |  | D1 | 0.171 | 0.246 | 0.334 |
|  |  | D2 | 0.18 | 0.061 | 0.16 |
|  | Norepinephrine | NET | -0.003 | 0.497 | 0.497 |
|  | Serotonin | 5-HTT | 0.013 | 0.479 | 0.497 |
|  |  | 5-HT1b | 0.165 | 0.142 | 0.273 |
|  |  | 5-HT6 | 0.276 | 0.003 | 0.055 |
|  |  | 5-HT2a | 0.313 | 0.024 | 0.083 |
|  |  | 5-HT4 | 0.316 | 0.034 | 0.103 |
|  |  | 5-HT1a | 0.408 | 0.019 | 0.083 |
|  | Acetylcholine | VAChT | 0.113 | 0.254 | 0.334 |
|  |  | M1 | 0.214 | 0.088 | 0.205 |
|  |  | α4β2 | 0.221 | 0.151 | 0.273 |
|  | Glutamate | NMDA | 0.065 | 0.323 | 0.377 |
|  |  | mGluR5 | 0.367 | 0.007 | 0.055 |
|  | GABA | GABA-A | -0.15 | 0.156 | 0.273 |
|  | Histamine | H3 | 0.209 | 0.192 | 0.31 |
|  | Cannabinoid | CB1 | 0.394 | 0.008 | 0.055 |
|  | Opioid | KOR | 0.156 | 0.212 | 0.318 |
|  |  | MOR | 0.396 | 0.022 | 0.083 |
|  | TSPO | TSPO | 0.127 | 0.283 | 0.35 |
| Cognition | Dopamine | D1 | 0.093 | 0.279 | 0.367 |
|  |  | D2 | 0.155 | 0.113 | 0.341 |
|  |  | DAT | 0.196 | 0.141 | 0.341 |
|  | Norepinephrine | NET | 0.283 | 0.018 | 0.125 |
|  | Serotonin | 5-HT2a | 0.003 | 0.471 | 0.471 |
|  |  | 5-HT1b | -0.049 | 0.314 | 0.375 |
|  |  | 5-HT4 | 0.07 | 0.424 | 0.445 |
|  |  | 5-HT6 | 0.074 | 0.408 | 0.445 |
|  |  | 5-HTT | 0.184 | 0.196 | 0.341 |
|  |  | 5-HT1a | 0.189 | 0.187 | 0.341 |
|  | Acetylcholine | M1 | -0.123 | 0.268 | 0.367 |
|  |  | α4β2 | 0.134 | 0.227 | 0.341 |
|  |  | VAChT* | 0.48* | 0.001* | 0.025* |
|  | Glutamate | NMDA | 0.1 | 0.211 | 0.341 |
|  |  | mGluR5 | 0.131 | 0.216 | 0.341 |
|  | GABA | GABA-A | -0.147 | 0.083 | 0.341 |
|  | Histamine | H3 | 0.271 | 0.032 | 0.167 |
|  | Cannabinoid | CB1 | 0.082 | 0.321 | 0.375 |
|  | Opioid | MOR | 0.196 | 0.145 | 0.341 |
|  |  | KOR | 0.414 | 0.006 | 0.063 |
|  | TSPO | TSPO | 0.201 | 0.204 | 0.341 |

***Note:*** *Spearman spatial correlations (ρ) between each significant PLSc-derived brain signature and all 21 receptor/transporter density maps, with spin-permutation p-values (p_spin_) and FDR-corrected q-values. Results are shown for both RSFC and anatomical signatures across all dimensions yielding a significant latent component. Associations surviving FDR correction (q<.05) are indicated with an asterisk.*

**Table S3. Full receptor and transporter annotation results across all dimensions in the anatomical model.**

| **Dimension** | **System** | **Receptor/Transporter** | **ρ** | **p_spin** | **q_FDR** |
| --- | --- | --- | --- | --- | --- |
| Positive  Symptoms | Dopamine | D1 | -0.092 | 0.361 | 0.401 |
|  |  | D2 | 0.207 | 0.152 | 0.341 |
|  |  | DAT | 0.253 | 0.099 | 0.313 |
|  | Norepinephrine | NET | -0.146 | 0.119 | 0.313 |
|  | Serotonin | 5-HT2a | -0.04 | 0.487 | 0.487 |
|  |  | 5-HT4 | 0.073 | 0.298 | 0.4 |
|  |  | 5-HT6 | -0.104 | 0.26 | 0.4 |
|  |  | 5-HT1a | 0.182 | 0.111 | 0.313 |
|  |  | 5-HTT | 0.248 | 0.051 | 0.313 |
|  |  | 5-HT1b | 0.281 | 0.055 | 0.313 |
|  | Acetylcholine | VAChT | -0.004 | 0.363 | 0.401 |
|  |  | M1 | -0.093 | 0.118 | 0.313 |
|  |  | α4β2 | -0.301 | 0.028 | 0.313 |
|  | Glutamate | NMDA | -0.017 | 0.435 | 0.457 |
|  |  | mGluR5 | -0.079 | 0.319 | 0.4 |
|  | GABA | GABA-A | -0.081 | 0.324 | 0.4 |
|  | Histamine | H3 | -0.137 | 0.185 | 0.354 |
|  | Cannabinoid | CB1 | -0.083 | 0.315 | 0.4 |
|  | Opioid | KOR | 0.094 | 0.318 | 0.4 |
|  |  | MOR | -0.151 | 0.162 | 0.341 |
|  | TSPO | TSPO | 0.216 | 0.077 | 0.313 |
| General | Dopamine | DAT | -0.081 | 0.356 | 0.447 |
|  |  | D1 | -0.35 | 0.103 | 0.269 |
|  |  | D2 | 0.459 | 0.011 | 0.085 |
|  | Norepinephrine | NET | 0.406 | 0.008 | 0.085 |
|  | Serotonin | 5-HTT | 0.013 | 0.467 | 0.482 |
|  |  | 5-HT1b | 0.026 | 0.467 | 0.482 |
|  |  | 5-HT6 | -0.137 | 0.362 | 0.447 |
|  |  | 5-HT2a | 0.175 | 0.225 | 0.41 |
|  |  | 5-HT4 | 0.325 | 0.05 | 0.175 |
|  |  | 5-HT1a | -0.417 | 0.035 | 0.146 |
|  | Acetylcholine | VAChT | -0.025 | 0.459 | 0.482 |
|  |  | M1 | 0.149 | 0.316 | 0.442 |
|  |  | α4β2 | 0.347 | 0.023 | 0.122 |
|  | Glutamate | NMDA | 0.123 | 0.287 | 0.442 |
|  |  | mGluR5 | 0.243 | 0.125 | 0.292 |
|  | GABA | GABA-A | 0.326 | 0.012 | 0.085 |
|  | Histamine | H3 | -0.17 | 0.296 | 0.442 |
|  | Cannabinoid | CB1 | 0.197 | 0.234 | 0.41 |
|  | Opioid | KOR | 0.016 | 0.482 | 0.482 |
|  |  | MOR | -0.224 | 0.232 | 0.41 |
|  | TSPO | TSPO | -0.397 | 0.093 | 0.269 |
| Cognition | Dopamine | D1 | -0.076 | 0.362 | 0.462 |
|  |  | D2 | 0.123 | 0.185 | 0.436 |
|  |  | DAT | 0.228 | 0.127 | 0.436 |
|  | Norepinephrine | NET | 0.179 | 0.116 | 0.436 |
|  | Serotonin | 5-HT2a | -0.023 | 0.477 | 0.497 |
|  |  | 5-HT1b | 0.067 | 0.374 | 0.462 |
|  |  | 5-HT4 | -0.086 | 0.27 | 0.436 |
|  |  | 5-HT6 | 0.115 | 0.26 | 0.436 |
|  |  | 5-HTT | 0.164 | 0.157 | 0.436 |
|  |  | 5-HT1a | -0.197 | 0.139 | 0.436 |
|  | Acetylcholine | M1 | 0.027 | 0.398 | 0.462 |
|  |  | α4β2 | 0.079 | 0.374 | 0.462 |
|  |  | VAChT* | 0.1 | 0.243 | 0.436 |
|  | Glutamate | NMDA | 0.141 | 0.186 | 0.436 |
|  |  | mGluR5 | 0.292 | 0.046 | 0.436 |
|  | GABA | GABA-A | 0.102 | 0.255 | 0.436 |
|  | Histamine | H3 | -0.161 | 0.24 | 0.436 |
|  | Cannabinoid | CB1 | 0.169 | 0.175 | 0.436 |
|  | Opioid | MOR | 0.044 | 0.418 | 0.462 |
|  |  | KOR | -0.095 | 0.334 | 0.462 |
|  | TSPO | TSPO | 0.015 | 0.497 | 0.497 |

***Note:*** *Spearman spatial correlations (ρ) between each significant PLSc-derived brain signature and all 21 receptor/transporter density maps, with spin-permutation p-values (p_spin_) and FDR-corrected q-values. Results are shown for both RSFC and anatomical signatures across all dimensions yielding a significant latent component. Associations surviving FDR correction (q<.05) are indicated with an asterisk.*
